# Lysosomal nucleotide metabolism regulates ER proteostasis through mTOR signaling

**DOI:** 10.1101/2020.04.18.048561

**Authors:** Keng Hou Mak, Qian Zhao, Pei-Wen Hu, Chi-Lam Au-Yeung, Jin Yang, Lita Duraine, Yong Yu, Monther Abu-Remaileh, David Sabatini, Jue D. Wang, Meng C. Wang

**Author notes:** Correspondence and requests for materials should be addressed to Meng C. Wang or Jue D. Wang.

## Abstract

Organelles in eukaryotic cells are compartmentalized to carry out different functions, and require specific mechanisms governing their coordination. Two of these organelles, the lysosome and the endoplasmic reticulum (ER), play crucial roles in regulating cellular homeostasis and organismal health. The lysosome contains various enzymes devoted to the hydrolysis of specific substrates, and dysfunctions of these hydrolases and their related metabolic processes are implicated in many diseases ^1^. On the other hand, ER is essential for protein synthesis and utilizes quality control mechanisms to maintain proteostasis. The metabolic status of the lysosome is now known to actively influence nuclear transcription and mitochondrial signaling ^2–4^. However, whether and how mechanistically lysosomal metabolic activities regulate ER proteostasis remain unclear. Here, we reported that RSH-1, the *Caenorhabditis elegans* RelA/SpoT Homolog (RSH) protein, carries a lysosomal NADPH phosphatase activity and acts through the mammalian/mechanistic target of rapamycin (mTOR) signaling to regulate ER proteostasis. We discovered that RSH-1 is localized to the lysosome. Its mutation reduces NADPH hydrolysis by the lysosome, leading to a protection against ER stress-induced lethality. We further revealed that this ER stress tolerance requires lysosomal vacuolar-type H^+^-ATPase (v-ATPase), and Rag GTPases and ribosomal protein S6 kinase (S6K) in the mTOR signaling pathway. Through transcriptome analysis, we discovered that the S6K activation by the RSH-1 mutation increases the basal expression of X-box binding protein 1 (XBP-1) and peptidyl-prolyl cis-trans isomerases, which is necessary for improving ER proteostasis. Moreover, we demonstrated the lysosomal localization of RSH-1 mammalian homolog and its conserved role in regulating mTOR signaling. Together, our findings reveal the key role of a specific lysosomal nucleotide hydrolase in regulating organellar coordination and also signaling mechanisms underlying this active regulation, and suggest that diseases resulted from deficiency in lysosomal hydrolases may be attributed to the distortion of signal transduction and organelle homeostasis.

---

RSHs are a superfamily of proteins that are well conserved across evolution in bacteria, plants, and metazoa ^5^, and regulate growth and stress responses. The RSH family of proteins in bacteria and plants carries both synthetase and hydrolase domains (Fig. 1a) that are known to synthesize and hydrolyze (p)ppGpp, a GDP/GTP-derived nucleotide messenger, respectively ^6–9^. On the other hand, the metazoan RSH carries only a hydrolase domain (Fig. 1a), and can dephosphorylate ppGpp and NADPH *in vitro* ^8,10^. *Drosophila* RSH homolog regulates starvation response ^8^, while the human homolog facilitates ferroptosis ^10^. In plants, RSH catalyzes (p)ppGpp metabolism within a well-defined localization in the chloroplast, an eukaryotic organelle with a bacterial origin ^7,9,11^. We thus hypothesized that the metazoan RSH might also have a specialized subcellular compartment, which would be mitochondria, another bacterium-originated organelle. However, to our surprise, we found that the *C. elegans* RSH-1(ZK909.3) protein is localized at lysosomes but not mitochondria (Fig. 1b, Extended Data Fig. 1a). In a transgenic strain carrying a C-terminal mRFP-tagged RSH-1 (RSH-1::mRFP), the fusion proteins are detected in filaments and puncta that overlap with the GFP fusions of the lysosomal membrane protein LMP-1 (Fig. 1b). The RSH-1::mRFP positive puncta are also co-localized with LysoTracker Green (Fig. 1b). On the other hand, we only observed a partial overlap between RSH-1::mRFP and RAB-7::GFP (Fig. 1b) that marks late endosomes ^12^, and there is no overlap between RSH-1::mRFP and mito-GFP labeled mitochondria (Extended Data Fig. 1a). We have also made another transgenic strain carrying a C-terminal Venus-tagged RSH-1 (RSH-1::Venus), and the fusion proteins are only detected at lysosomes upon increased lysosomal pH by v-ATPase inactivation (Extended Data Fig. 1b). Given that the pKa (the pH at which fluorescence intensity reduces by half) of mRFP and Venus is 4.5 and 6.0, respectively ^13^, these results suggest that RSH-1 fusion proteins are likely localized to the acidic lumen of the lysosome (pH 4.5~5.5) ^14^.

**Figure 1.**
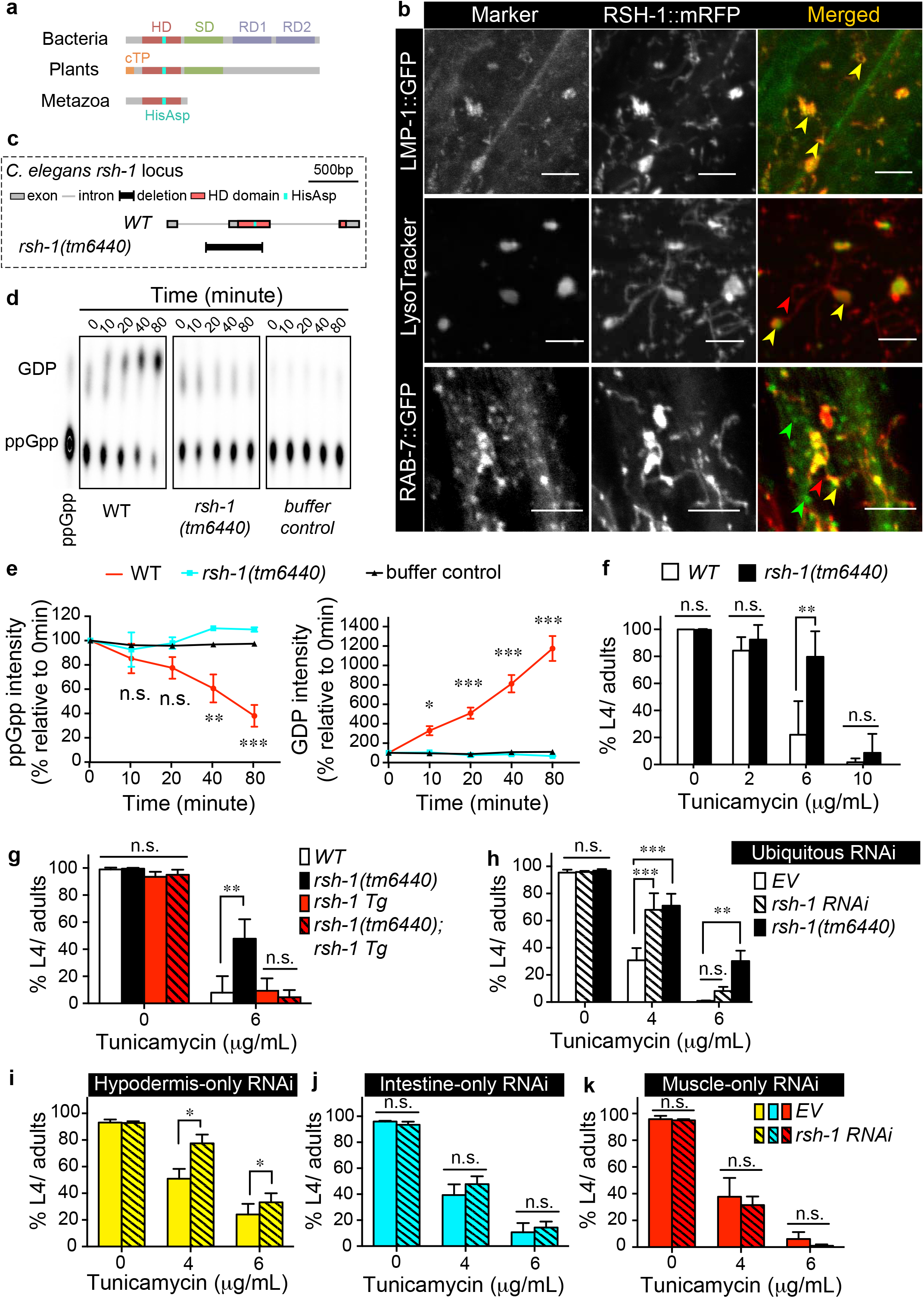
Lysosomal RSH-1 regulates ER proteostasis. **a**) Diagram of RSH conserved domains across different kingdoms. HD, hydrolase domain. SD, synthetase domain. RD, regulatory domain. cTP, chloroplast targeting peptide. His-Asp, conserved catalytic residues. **b**) Funtional RSH-1::mRFP fusion proteins co-localize with lysosomal markers (LMP-1::GFP and LysoTracker green DND-26) in hypodermal cells. It overlaps partially with the late endosomal marker RAB-7::GFP. mRFP is fused at the C terminus of RSH-1 and the fusion protein fully rescues the phenotype of the *rsh-1(tm6440)* mutants (**g**). Yellow, green and red arrows indicate overlapping signals, RAB-7::GFP positive signals alone, and RSH-1::mRFP positive signals alone, respectively. Scale bar = 5μm. **c**) The deletion allele *tm6440* disrupts the hydrolase domain of RSH-1 and removes the conserved catalytic His-Asp residues. **d**) Worm lysate from wild type (WT) but not the *rsh-1(tm6440)* mutants can hydrolyze P^32^-labled ppGpp. After incubation with whole worm lysate, P^32^-labled ppGpp and their GDP products are detected on 2D thin-layer chromatography. **e**)Quantification of data from (**d**) and two other independent replicates. Signal intensities are relative to 0 min of the same samples. **f**) The *rsh-1(tm6440)* mutants are more resistant to tunicamycin-induced ER stress than WT (*n*=3 replicates). **g**) Transgenic restoration of *rsh-1* expression using its endogenous promoter (*rsh-1 Tg*) suppresses the resistance of the *rsh-1(tm6440)* mutants to tunicamycin-induced ER stress (*n* = 4 replicates). **h**-**k**) Ubiquitous RNAi inactivation of *rsh-1* increases tolerance to tunicamycin-induced ER stress (*n* = 5 replicates for empty vehicle (*EV*), *n* = 4 replicates for *rsh-1 RNAi* and *rsh-1(tm6440)*). Hypodermis (**i**) - but not intestine (**j**) - or muscle (**k**)-specific inactivation of *rsh-1* by RNAi is sufficient to induce ER tolerance against tunicamycin (*n* = 4 replicates for each condition). Error bars represent SEM. \*\*\**p<0.001*, \*\**p<0.01, *p<0.05*, n.s. *p>0.05* by two-way ANOVA.

*C. elegans* RSH-1 contains two conserved catalytic residues His113 and Asp114 in the hydrolase domain, which are lacking in the *rsh-1*(*tm6440)* mutant (Fig. 1c). To test whether this mutation of *rsh-1* disrupts its enzymatic activity, we measured the hydrolytic ability of worm lysates toward ppGpp. We found that the protein lysate from wild type, but not the *rsh-1*(*tm6440)* mutant can dephosphorylate ppGpp into GDP (Fig. 1d and e), suggesting that the phosphatase activity is absent in the *rsh-1*(*tm6440)* mutant. Using this loss-of-function mutant, we next examined whether the loss of lysosomal RSH-1 affects animal physiology. We found that under normal conditions, the *rsh-1(tm1440)* mutants have slightly delayed development and reduced brood size but are otherwise phenotypically wild type in their body length and food intake rates (Extended Data Fig. 2a-e). Upon starvation or heat shock, the *rsh-1* mutants show no difference in their survival rates, when compared to wild type (Extended Data Fig. 2f, 2g). On the other hand, the *rsh-1* mutants show increased sensitivity to oxidative stress induced by hydrogen peroxide (Extended Data Fig. 2h) but have enhanced tolerance to cold shock (Extended Data Fig. 2i) and ER stress induced by either tunicamycin (Fig. 1f) or dithiothreitol (Extended Data Fig. 2j). Together, these results reveal the significance and specificity of *rsh-1* in regulating different stress responses. In particular, given that ER only exists in eukaryotic cells, the regulation of ER stress tolerance is a unique function of RSH-1 distinct from its prokaryotic homologs.

To confirm this ER regulation by RSH-1, we first restored the expression of *rsh-1* using its endogenous promoter and showed that it can suppress the enhanced ER stress tolerance in the mutants (Fig. 1g). We also knocked down *rsh-1* using RNA interference (RNAi) and observed enhanced ER stress tolerance (Fig. 1h). To further investigate the key tissue for RSH-1-mediated ER regulation, we generated a transgenic *C. elegans* strain carrying a GFP reporter driven by the *rsh-1* genomic regulatory elements, including the 1.7kb upstream promoter and the full open reading frame. We found that *rsh-1* predominantly expresses in hypodermis, muscle and intestine (Extended Data Fig. 3a), which are the tissues actively participating in metabolic and stress responses. We then employed tissue-specific RNAi to knockdown *rsh-1* and examined their effects on ER stress tolerance. We found that RNAi inactivation of *rsh-1* only in hypodermal cells is sufficient to enhance the tolerance against ER stress (Fig. 1i), but intestine- or muscle-specific inactivation of *rsh-1* has no such effects (Fig. 1j and k). Together with the lysosomal localization of RSH-1 in the hypodermis (Fig. 1b), these results suggest a cell-autonomous regulation of ER proteostasis by lysosomal RSH-1.

To understand how lysosomal RSH-1 regulates ER proteostasis, we first examined whether lysosomal activity is altered in the *rsh-1* mutant. Using LysoTracker DND-26 staining, we measured the number of mature acidic lysosomes and found that the *rsh-1* mutant shows a decreased number in hypodermal cells (Fig. 2a). Furthermore, transmission electron microscopy (TEM) reveals that in the *rsh-1* mutants, the numbers of multi-lamellar bodies are increased (Fig. 2b), which is likely due to inefficient lysosomal degradation. Together, these results suggest that the *rsh-1* mutation reduces lysosomal activity. To test whether the reduction in lysosomal activity is responsible for the increased ER stress tolerance, we knocked down 142 genes involved in lysosomal functions by RNAi and screened their effects on ER stress response (Supplementary Table 1). We found that RNAi inactivation of v-ATPase components decreases ER stress tolerance (Supplementary Table 1), suggesting that reducing lysosomal activity is not sufficient to increase ER stress tolerance. More importantly, we identified that RNAi inactivation of *vha-17*, encoding the *e* subunit of the membrane-bound V_0_ domain of v-ATPase, specifically abrogates the ER stress tolerance in the *rsh-1* mutants (Fig. 2c). In contrast, RNAi inactivation of other components of v-ATPase, for example *vha-8* encoding the E subunit of the cytoplasmic V_1_ domain of v-ATPase, has no such effect (Fig. 2d, Supplementary Table 1). Together, these results suggest that RSH-1-mediated regulation of ER proteostasis is not simply a result of lysosomal dysfunction, but act through specific lysosomal v-ATPase components.

**Figure 2.**
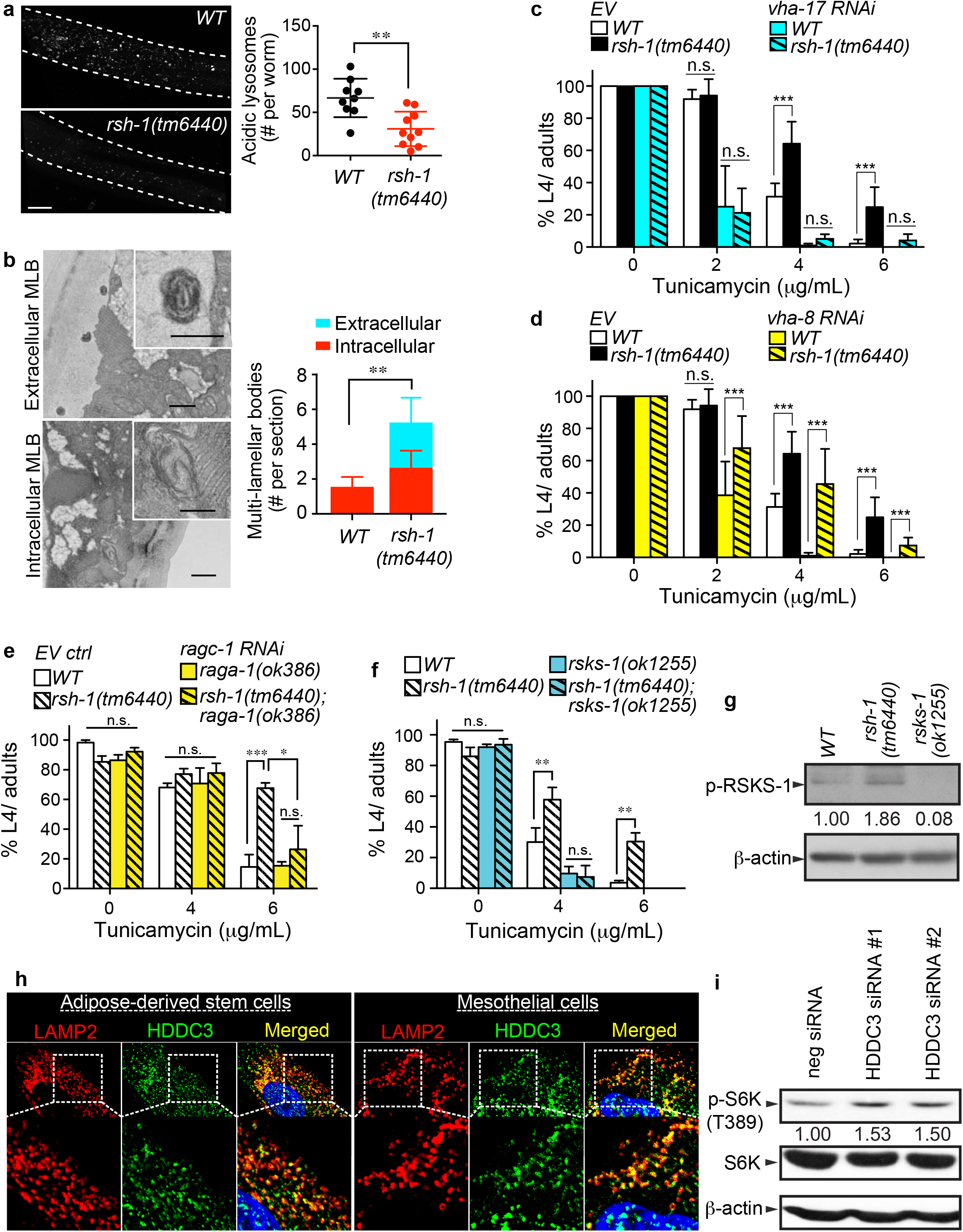
RSH-1 regulates ER proteostasis via mTOR signaling. **a**) The *rsh-1(tm6440)* mutants show less mature acidic lysosomes (marked by LysoTracker Green DND-26) in hypodermis under an unstressed condition. Scale bar = 20μm. Unpaired *t-*test (two-way). *n* = 9 worms for *WT*, *n* = 10 worms for *rsh-1(tm6440)*. Error bars represent mean+/− SD. **b**) Accumulation of multi-lamellar bodies (MLB) in the *rsh-1(tm6440)* mutants. Representative TEM images show extracellular and intracellular MLB in the *rsh-1(tm6440)* mutants, which are either not detected or less in WT. Scale bar = 400nm (Inset: 200nm). Error bars represent SEM. *n* = 6 hypodermal sections for *WT*, *n* = 5 hypodermal sections for *rsh-1(tm6440)*. **c** and **d**) RNAi inactivation of a v-ATPase component, *vha-17* (**c**) or *vha-8* (**d**), increases ER stress sensitivity on its own, but only *vha-17* RNAi inactivation suppresses the ER stress tolerance in the *rsh-1(tm6440)* mutants (**c**). Error bars represent SEM. *n* = 5 replicates for each genotype. **e**) Inactivation of Ragulator components in mTOR signaling, RAGA/*raga-1(ok386)* and RAGC/*ragc-1(RNAi)*, suppresses the protective effect of the *rsh-1* mutation against ER stress. Error bars represent SEM. *n* = 4 replicates for *WT* and *rsh-1(tm6440), n*=5 replicates for *raga-1(ok386)*; *ragc-1(RNAi)* and *rsh-1(tm6440); raga-1(ok386)*; *ragc-1(RNAi)*. **f**) The *rsks-1(ok1255)* deletion decreases ER stress tolerance and also suppresses the tolerance in the *rsh-1(tm6440)* mutants. Error bars represent SEM. *n*=4 replicates for *WT* and *rsh-1(tm6440)*, *n*=3 replicates for the *rsks-1(ok1255)* and *rsh-1(tm6440); rsks-1(ok1255)*. **g**) The level of phosphorylated RSKS-1(Thr404) is increased in the *rsh-1(tm6440)* mutants under an unstressed condition. The values indicate relative levels of phosphorylated RSKS-1 after normalization to β-actin. An independent replicate is shown in Extended Data Fig. 4a. **h**) HDDC3, encoded by the human homolog of RSH-1, colocalizes with lysosomal marker LAMP2 in human adipose-derived stem cells and mesothelial cells. **i**) The level of phosphorylated p70 S6K is increased in HeLa cells after transfection with HDDC3-specific siRNAs, compared to control siRNA. Relative p-S6K levels normalized to the corresponding total p70 S6 kinase level are indicated. Independent replicates are shown in Extended Data Fig. 4d. \*\*\**p<0.001*, \*\**p<0.01, *p<0.05*, n.s. *p>0.05* by *t-*test (a, b) or two-way ANOVA (c-f).

Lysosomal v-ATPase complex is not only required for lysosomal acidification, but also mediates the lysosomal activation of mTOR through cooperating with the Rag GTPases and Ragulator complex ^15,16^. In *C. elegans, raga-1* and *ragc-1* encode the RagA and RagC GTPase homologs, respectively. We found that inactivation of both *raga-1* and *ragc-1* does not affect ER stress response, but specifically suppresses the ER stress tolerance in the *rsh-1* mutants (Fig. 2e). These results suggest that the *rsh-1* mutation requires lysosomal activation of mTOR to regulate ER proteostasis. We next assessed RSKS-1, the *C. elegans* homolog of S6K that is a key effector phosphorylated by mTOR at Thr404 (Thr389 in human) ^17,18^and expresses predominantly in the hypodermis ^19^ where *rsh-1* acts to regulate ER stress response. We found that the *rsks-1* mutation fully abrogates the ER stress tolerance in the *rsh-1* mutants (Fig. 2f), and the phosphorylation of RSKS-1 is increased in the *rsh-1* mutants (Fig. 2g, Extended Data Fig. 4a).

Next, we studied the molecular mechanism by which mTOR activation improves ER proteostasis in response to RSH-1 deficiency. Through RNA-seq transcriptome analyses, we identified 560 genes that are differentially expressed in the *rsh-1* mutants compared to wild type (Fig. 3a). For 558 of these genes, their expression changes are fully dependent on *rsks-1* (Fig. 3b), suggesting RSKS-1 as a master regulator of the transcriptional responses downstream of RSH-1. Among those candidate genes, we found that *xbp-1* is up-regulated in the *rsh-1* mutants, which is dependent on *rsks-1* (Fig. 3c). XBP-1 is a transcription factor involved in ER unfolded protein response (UPR^ER^), which is well conserved from yeast to mammals, and its activation is regulated post-transcriptionally by non-conventional splicing of an intron from its transcript ^20,21^. The spliced form of XBP-1 can then function as an active transcription factor to induce ER chaperones, which increase the protein-folding capacity at ER ^20^. We found that the level of the spliced active form of *xbp-1* is also up-regulated in the *rsh-1* mutants (Fig. 3c); while upon ER stress, this level is increased in wild type but not further enhanced in the *rsh-1* mutants (Fig. 3d). As a key transcription factor for UPR^ER^, XBP-1 mediates the induction of *hsp-4*, the *C. elegans* homolog of mammalian ER chaperone BiP, which can be visualized in both the hypodermis and the intestine using a *hsp-4* GFP reporter ^20,22^. We found that upon ER stress, the induction level of *hsp-4* in the hypodermis is significantly higher in the *rsh-1* mutants than that in wild type (Fig. 3e, *p<0.001*, Extended Data Fig. 3b). On the other hand, the induction level of *hsp-4* in the intestine is similar between the mutants and wild type (Fig. 3e, Extended Data Fig. 3b). This elevated induction of BiP chaperones can enhance protein-folding capacity, accelerate ER stress resolution, and ultimately contribute to ER stress tolerance. Thus, a primed activation of XBP-1 occurs in the *rsh-1* mutants, which can increase ER proteostasis and promote a better resolution of ER stress. In supporting this idea, we found that RNAi inactivation of *xbp-1* fully blocked the transcriptional induction of *hsp-4* (Fig. 3f) and the enhanced ER tolerance (Fig. 3g) in the *rsh-1* mutants.

**Figure 3.**
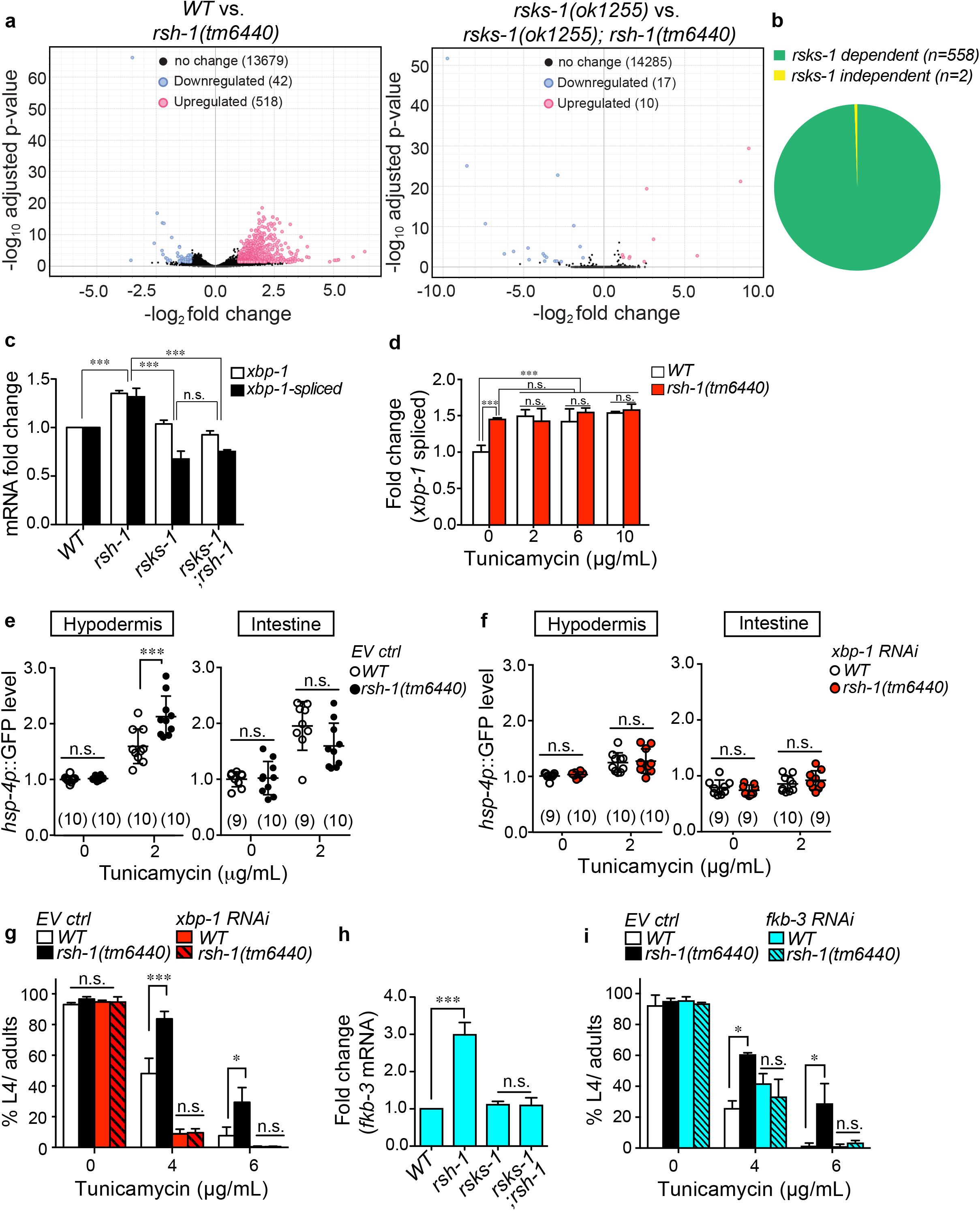
RSH-1 regulates ER proteostasis through mTOR-mediated transcriptional responses. **a**) RNA-seq analysis reveals the transcriptional difference between WT and the *rsh-1(tm6440)* mutants (left panel), and between the *rsks-1(ok1255)* mutants and the *rsks-1(ok1255);rsh-1(tm6440)* double mutants (right panel). A two-fold change and a false discovery rate (FDR) < 0.05 are used as thresholds to identify differentially expressed genes. **b**) 99.6 % of the differentially expressed genes in the *rsh-1(tm6440)* mutants are dependent on *rsks-1.* **c**) qRT-PCR assesses the levels of the unspliced and the spliced (activated) forms of *xbp-1* transcripts. In an unstressed condition, the *rsh-1(tm6440)* mutants show higher levels of both forms than WT. The *rsks-1(ok1255)* deletion decreases the level of active spliced *xbp-1* and abolishes the induction of both forms in the *rsh-1(tm6440)* mutants. Error bars represent SEM. *n* = 6 replicates for WT and *rsh-1(tm6440)*, *n* = 3 replicates for *rsks-1(ok1255)* and *rsks-1(ok1255); rsh-1(tm6440)*. **d**) qRT-PCR assesses the level of the active spliced form of *xbp-1* transcripts. The tunicamycin treatment induces the level of active spliced *xbp-1* in WT but does not further enhance its induction in the *rsh-1(tm6440)* mutants. Error bars represent SEM. *n* = 6 replicates for WT and *rsh-1(tm6440)*. **e** and **f**) Upon the tunicamycin treatment, the level of the *hsp-4* (ER chaperone) GFP reporter is increased in both hypodermal and intestinal cells, and the *hsp-4* induction in hypodermal cells is specifically enhanced in the *rsh-1(tm6440)* mutants compared to WT (**e**). The enhanced hypodermal induction of *hsp-4* is fully abrogated by the RNAi inactivation of *xbp-1* (**f**). Error bars represent SD. Sample size (n) shown in figure panels. **g**) RNAi inactivation of *xbp-1* completely suppresses the ER stress resistance conferred by the *rsh-1(tm6440)* mutation. Error bars represent SEM. *n* = 5 replicates for each genotype. **h**) qRT-PCR assesses the level of *fkb-3*, encoding a peptidyl-prolyl cis-trans isomerase. The *rsh-1(tm6440)* mutants show a higher level than WT, which is suppressed by the *rsks-1(ok1255)* deletion. Error bars represent SEM. *n* =3 for all genotypes. **i**) RNAi inactivation of *fkb-3* completely suppresses the ER stress resistance conferred by the *rsh-1(tm6440)* mutation. Error bars represent SEM. *n* = 5 replicates for each genotype. \*\*\**p<0.001, *p<0.05*, n.s. *p>0.05* by two-way ANOVA (c-g, and i) or *t*-test (h).

In addition, we identified a class of genes *fkb-3,4,5,7,8* that are up-regulated in the *rsh-1* mutants in a *rsks-1* dependent manner (Fig. 3h, Extended Data Fig. 4b). These genes encode peptidyl-prolyl cis-trans isomerases, which catalyze the cis-trans isomerization of proline imidic peptide bonds and facilitate protein folding ^23^, and FKB-3, 4, 5 contain ER signal peptides (Extended Data Fig. 4c). We found that RNAi inactivation of *fkb-3* fully suppresses the ER stress tolerance in the *rsh-1* mutants (Fig. 3i). Taken together, these results suggest that lysosomal RSH-1 regulates ER proteostasis through mTOR-mediated transcriptional responses.

Furthermore, we found that the function of lysosomal RSH-1 in regulating mTOR activity is conserved in mammalian cells. The human homolog of RSH-1, HDDC3 co-localizes with LAMP2, the lysosomal protein marker in human adipose-derived stem cells and mesothelial cells (Fig. 2h), and its RNAi knockdown enhances the phosphorylation of S6K (Fig. 2i, Extended Data Fig. 4d). Proteome-based analysis of macromolecular complexes has revealed the interaction between RSH and v-ATPase in four different metazoan species, including HDDC3 and ATP6V1C1 in human, Mesh1 and Vha44 in *Drosophila*, RSH-1 and VHA-11 in *C. elegans*, and SPU_016827 and Sp_Atp6v1c1 in sea urchin ^24^. Thus, metazoan RSH likely acts through lysosomal v-ATPase to regulate mTOR activation.

Next, to characterize the metabolic activity of RSH-1 and its roles in regulating mTOR activity and ER proteostasis, we conducted RNAi screens of genes that are involved in nucleotide biosynthesis and degradation (supplementary table 2). We found that RNAi inactivation of R151.2, *nadk-1* and *nadk-2*, encoding the *C. elegans* homologs of phosphoribosyl pyrophosphate synthetases (PRPS) and NAD kinases (NADK), respectively, fully suppress the ER stress tolerance in the *rsh-1* mutants (Fig. 4a-c). PRPS and NADK can be both linked to NADPH biosynthesis (Fig. 4d), and their involvements suggest an alteration of NADPH metabolism in the *rsh-1* mutants. Consistently, a recent study shows that human RSH homolog, MESH1 dephosphorylates NADPH ^10^. To examine whether lysosomes carry a previously unknown NADPH phosphatase activity catalyzed by RSH-1, we immunopurified lysosomes from the *rsh-1* mutant and wild type worms, incubated purified lysosomes with NADPH and measured the production of free phosphates. We found that purified lysosomes from wild type worms are sufficient to dephosphorylate NADPH, but this NADPH dephosphorylation activity is decreased when using lysosomes purified from the *rsh-1* mutants (Fig. 4e). Furthermore. this NADPH phosphatase activity is dependent on manganese ion (Mn^2+^) (Fig. 4e,f), a cofactor also required for bacterial (p)ppGpp hydrolase ^25^. We also conducted structural simulation of RSH-1 with NADPH as a substrate based on the crystal structure of its *Drosophila* and human homologs ^10^. Structural analysis indicates that residues for the molecular recognition of NADPH are highly conserved (Fig. 4g). Together, these results suggest that RSH-1 contributes to Mn^2+^-dependent NADPH hydrolysis at the lysosome.

**Figure 4.**
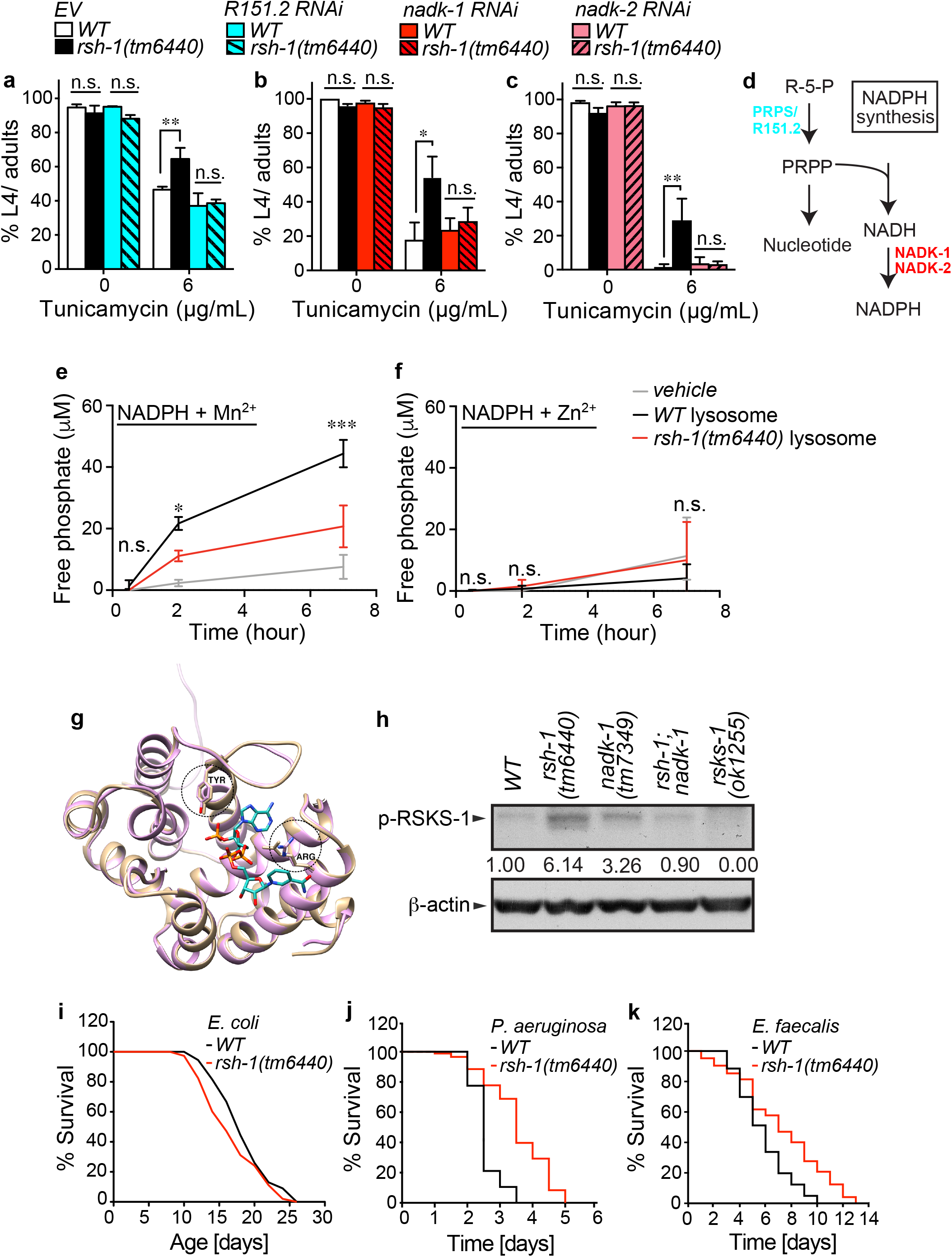
RSH-1 controls NADPH metabolism to regulate mTOR and ER. **a**-**d)** RNAi inactivation of *R151.2* (**a**), *nadk-1* (**b**) or *nadk-2* (**c**) fully suppresses the ER stress tolerance in the *rsh-1(tm6440)* mutants. R151.2, NADK-1 and NADK-2 are involved in NADPH metabolism (**d**). Two-way ANOVA. *n* =3 replicates for a, b, *n* =4 replicates for c. **e** and **f**) Lysosomes purified from WT can dephosphorylate NADPH, and this activity is detected with the manganese ion (Mn^2+^, **e**), but not the zinc ion (Zn^2+^, **f**) buffer. Lysosomes purified from the *rsh-1(tm6440)* mutants show reduced activities in dephosphorylating NADPH. Two-way ANOVA comparing *WT* and the *rsh-1* mutant samples. Error bars represent standard deviations (SD). *n*= 3 replicates for each time point. **g**) Structure comparison of the NADPH binding site of human RSH structure (gold) with *C. elegans* RSH homology model (pink). The NADPH molecule and two conserved amino acids around NADPH are shown in a stick model. **h**) The *nadk-1(tm7349)* mutation suppresses the increased RSKS-1 phosphorylation in the *rsh-1(tm6440)* mutants, and also induces RSKS-1 phosphorylation on its own. Independent replicates are shown in Extended Data Fig. 4e. **i**) The lifespan of the *rsh-1(tm6440)* mutants is not significantly different from WT. n=100 for WT, n=110 for *rsh-1(tm6440)*. *p>0.05*, Log-rank test. **j** and **k**) The *rsh-1(tm6440)* mutants are more resistant to the pathogenic infection induced by *Pseudomonas aeruginosa* (**j**) or *Enterococcus faecalis* (**k**). *n* = 90 animals per genotype. *p<0.001, p<0.05* by Log-rank test.

We further confirmed that the *nadk-1* mutation suppresses the active phosphorylation of RSKS-1 in the *rsh-1* mutants (Fig. 4h, Extended Data Fig. 4e), suggesting that increased NADPH is responsible for the mTOR activation. Interestingly, the *nadk-1* mutation alone also induces RSKS-1 phosphorylation (Fig. 4h, Extended Data Fig. 4e), suggesting that mTOR activity is sensitive to the NADH/NADPH balance. We also examined *cup-5*, encoding the *C. elegans* orthologue of human TRPML1 calcium-release channel, which is localized at the lysosome ^26^. TRPML1 is a target of NAADP, a calcium-mobilizing nucleotide that is synthesized from NADPH ^27^, and mTOR can be activated by lysosomal calcium release through TRPML1 ^28^. We found that RNAi inactivation of *cup-5* does not affect the ER stress tolerance in the *rsh-1* mutant (Extended Data Fig. 4f), suggesting that RSH-1-mediated lysosomal NADPH hydrolysis unlikely acts through NAADP-sensitive calcium release to regulate ER proteostasis and mTOR activity.

Genetic reduction or pharmaceutical inhibition of mTOR signaling extends lifespan in diverse organisms from yeast to mice ^29^, and mutations in Rag GTPases that mediate the lysosomal activation of mTOR extend lifespan in *C. elegans* ^30–32^. However, when we measured the lifespan of the *rsh-1* mutants, we found no significant changes compared to wild type controls (Fig. 4i, p>0.05), suggesting that the mTOR activation by RSH-1 knockdown does not affect the survival of animals under normal conditions. On the other hand, ER proteostasis has been linked to the protection against pathogenic infection through XBP-1 ^33^. We next examined pathogenic resistance in the *rsh-1* mutants. We found that when exposed to pathogenic bacteria, *Pseudomonas aeruginosa* PA14 or *Enterococcus faecalis* OG1RF, the *rsh-1* mutants show an increased survival compared to wild type controls (Fig. 4j, k). Thus, through the induction of mTOR signaling and the consequent enhancement of ER proteostasis, lysosomal NADPH accumulation exerts a protective effect against pathogenic infection.

In summary, our findings demonstrate that lysosomal nucleotide metabolism can actively regulate ER proteostasis through specific signaling mechanisms (Fig. 5). RSH localized at the lysosome binds to v-ATPase and catalyzes the hydrolysis of NADPH, and upon RSH inactivation, NADPH accumulation leads to the activation of mTOR and S6K and consequently the improvement of ER proteostasis through the transcriptional upregulation of UPR^ER^ factors and protein folding machinery (Fig. 5).

**Figure 5.**
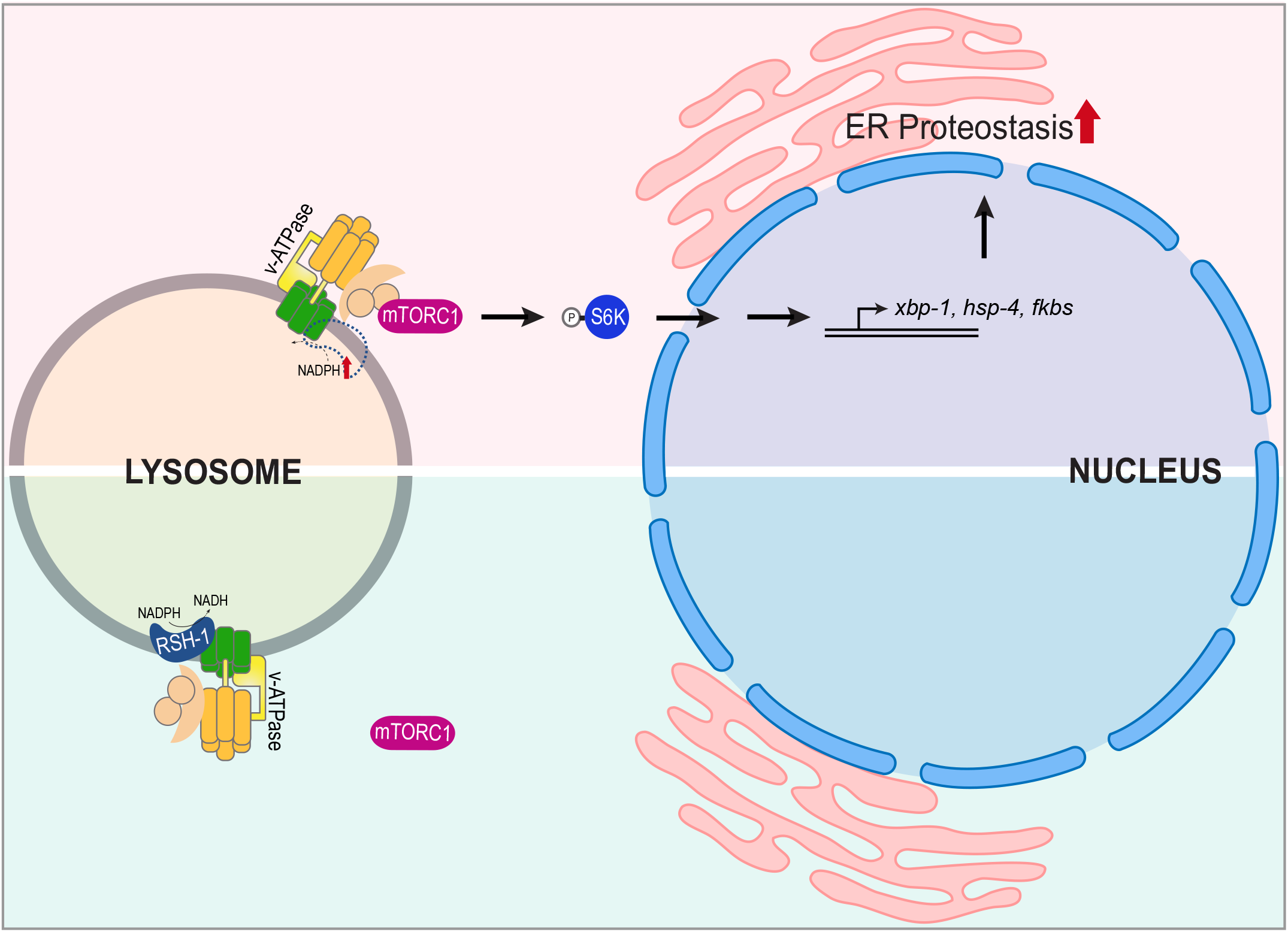
Schematic model of RSH-1 linking lysosome and ER homeostasis. RSH-1 is a lysosomal NADPH hydrolase associated with v-ATPase. Upon its inactivation, NADPH accumulation activates mTOR and S6K, which in turn enhances ER proteostasis through transcriptional induction of UPR^ER^ factors and protein folding enzymes.

NADPH is a crucial reducing agent involved in anabolic reactions. Our studies suggest that its induction at the lysosome can activate mTOR, the master regulator of anabolic processes, and at the same time enhance ER proteostasis to accommodate increased protein synthesis. Thus, in addition to lysosomal amino acids and cholesterol ^34,35^, nucleotide inputs can also drive mTOR activation through v-ATPase at the lysosome. This mechanism coordinates lysosomal nucleotide metabolism, lysosome-to-nucleus signal transduction, and ER homeostasis. In bacteria, RSH hydrolase activity can be regulated at the transcription level, as well as by allosteric mechanisms through binding with nutrient sensors ^36^. In particular, branched-chain amino acids can act through these allosteric mechanisms to inhibit the hydrolase activity toward ppGpp ^37^, and in metazoan cells, the branched-chain amino acid leucine is a potent activator of mTOR ^38–40^. Thus, it would be interesting for future studies to characterize whether and how amino acids and other metabolic cues could fine tune the activity of metazoan RSH hydrolases through allosteric mechanisms either directly (binding to RSH) or indirectly (binding to RSH interactors). Despite its phosphatase activity toward NADPH, RSH might catalyze other nucleotide targets in metazoan cells including (p)ppGpp that may be synthesized *de novo* by metazoan cells or taken up from bacteria. Future studies might reveal these nucleotide targets and their roles in regulating lysosomal signaling. Together, investigation of these unexplored regulatory mechanisms will advance our current understanding on organelle crosstalk, metabolic sensing and signaling, and stress and immune responses.

## Methods

### *C. elegans* strains and maintenance

The following strains were obtained from *Caenorhabditis* Genetics Center: RT258*(pwIs50[lmp-1::GFP + Cb-unc-119(+)];unc-119(ed3))*, DA2123*(adIs2122 [lgg-1p::GFP::lgg-1 + rol-6(su1006)])*, VC222*(raga-1(ok386))*, RB1206*(rsks-1(ok1255))*, and WM27*(rde-1(ne219))*. The following strains were gifts: XW5735*(ls[phyp-7::gfp::rab-7])* is from Xiaochen Wang lab; JM45*(rde-1 (ne219); Is[ges-1p::RDE-1::unc54 3’UTR, myo2p::RFP3])*, JM43*(rde-1 (ne219); Is[wrt-2p::RDE-1::unc54 3’UTR, myo2p::RFP3])*, and *rde-1 (ne219); Is[myo-3p::RDE-1::unc54 3’UTR, myo2p::RFP3]* are from J. Mello; and SJ4005*(zcIs4 [hsp-4::GFP])* is from J. Mamrosh. The strains *zk909.3(tm6440)* and *nadk-1(tm7349)* were obtained from National BioResource Project (Japan) and backcrossed to N2 seven times.

Strains were maintained at 20^°^C on NGM agar plates seeded with OP50 *E. coli*., or HT115 *E. coli* for RNAi experiments, and kept at least three generations without starvation before experiments.

### Transgenic strain construction

DNA construct for the transcriptional reporter *raxls27[rsh-1p::rsh-1::sl2::gfp; pmyo-2::mcherry]* or translational reporter *raxIs48[rsh-1p::rsh-1cDNA::mRFP]* was injected with a co-injection marker and salmon sperm DNA into young adult germline. Transgenic progenies with high transmission rate were gamma irradiated for extrachromosomal array integration and backcrossed to wildtype N2 (×3 for *raxIs27* and ×10 for *raxIs48*).

#### DNA construct for rsh-1 transcriptional reporter raxls27[rsh-1p::rsh-1::sl2::gfp]

The *rsh-1* genomic region, consisting of its 1.7kb promoter and ORF, was amplified from N2 worm lysate and fused to a *sl2::gfp* fragment by PCR fusion method ^41^ to create a polycistronic transgene. The A* primer is 5’-tgttcatacattttcttgcccc-3’ and the D primer is 5’-aaagtaggatgagacagcggtaccttaggccaaatgacggttgat-3’.

#### DNA construct for rsh-1 translational reporter raxIs48[rsh-1p::rsh-1cDNA::mRFP]

The *rsh-1* 1.7kb promoter and its cDNA were cloned into the restriction enzyme sites BamHI and SmaI, respectively, of pPD95.79 (mRFP) vector. The primers for cloning the promoter were 5’-gtctcgggatccatttcttcccttcctttctg-3’(FW) and 5’-gagagaggatcctgatttctagaaattaattatt-3’(RV), and for cloning the cDNA 5’-attattacccgggggccaaatgacggttgat-3’ (FW) and 5’-tgagagacccgggatggtcgacaaaaaaca-3’(RV).

#### DNA construct for rsh-1 translational reporter raxEx523[rsh-1p::rsh-1cDNA::Venus]

It was created by NEBuilder HiFi DNA Assembly. The fragment contains *rsh-1* 1.7kb promoter and its cDNA was amplified from previous construct for *rsh-1::mRFP* reporter with primer 5’-gctcggatccactagtaacgggtcgactctagaggatcca-3’ (FW) and 5’-ttactcattccagaacctccggccaaatgacggttgatg-3’(RV). The fragment contains Venus and *unc-54* 3’UTR was amplified from previous pCR2.1-TOPO (Venus) construct with primers 5’-ggaggttctggaatgagtaaaggagaagaacttttc-3’ (FW) and 5’-ttactagtggatccgagc-3’(RV). The construct was linearized by ApaI before microinjection.

### ppGpp hydrolysis assay with whole worm lysate

Synchronized L4 animals were collected, rinsed with PBS three times, and lysed by grinding with pellet pestle until no visible animals were seen under microscope. After removing debris with centrifugation, the lysate was snap frozen. To perform ppGpp hydroslysis assay, [α-^32^P]-ppGpp was synthesized *in vitro* as previously described ^42^ with these modifications listed: 1) 2 mM ATP and 0.25 mM [α-^32^P]-GTP was used in the 300 μL synthesis reaction mix; 2) the reaction mix was purified using anion exchange column (HiTrap QFF 1 mL; GE Healthcare); 3) 1 mL elution fraction with 0.5 M LiCl was collected as 2X [α-^32^P]-ppGpp stock solution. Then, [α-^32^P]-ppGpp was incubated with the lysate containing 10 μg of total protein in 50 μL buffer containing 50 mM Tris pH 8.0 and 200 mM NaCl. The reaction was stopped by adding formic acid to 333 mM final concentration at indicated time points. [α-^32^P]-ppGpp and [α-^32^P]-GDP in the reaction were detected and quantified using thin-layer chromatography as previously described ^43^.

### Lysosomal NADPH phosphatase activity assay

Immunopurification of lysosomes is modified based on the method used in mammalian cells ^44^. Briefly, we generated a transgenic strain expressing an extrachromosomal array of C-terminal RFP- and HA- tagged lysosomal membrane protein LMP-1 in hypodermal cells (*col-12p::lmp-1::rfp::3xha*). Around 50,000 L4 animals per genotype were collected, rinsed with PBS, ground with pellet pestle on ice until no visible animals were seen under microscope. The lysate was spun at 1,000g for 3 min in cold room to remove debris and then the supernatant was incubated with anti-HA magnetic beads (Thermo Fisher Scientific) for six minutes at room temperature. The bound fraction was washed four times with PBS. Equal amounts of bound fraction (3.4ug total protein based on BCA assay results) were then used for NADPH phosphatase assay. The assay was performed with 1mM NADPH in a buffer containing 50mM Tris pH7.5, 200mM NaCl and 1mM MnCl_2_ or ZnCl_2_. An aliquot was removed from the reaction at 0.5-, 2-, and 7-hour time points using a magnetic stand. The amounts of released phosphate were quantified with Malachite Green Phosphate Assay Kit (Cayman Chemical).

### Physiological measurements of *C. elegans*

#### Food intake

Adult individuals (86 hours post egg laying at 20°C) were placed on OP50-seeded NGM plates and their anterior regions were video-recorded for thirty seconds using stereoscope. The videos were played at slow motion to count numbers of pharyngeal contractions.

#### Developmental timing

Gravid adults are placed on OP50-seeded plates to lay eggs for an hour. Around thirty progenies were laid per genotype. After 60 hours of growth at 20°C, the progenies were scored every two hours for reaching late L4 (M-shaped vulva) or older age. The experiment was repeated three times.

#### Brood size measurement

Ten L4 hermaphrodite larvae were maintained at 20°C, and transferred daily to new, individual plates until reproduction stopped. Brood size for each individual was the total number of hatched progenies across the reproductive span. Animals lost during the experiment were not counted. The experiment was repeated three times.

#### Body Size

At least ten D1 (72 hours post egg laying) adults, grown on OP50 at 20°C, were imaged with stereoscope SMZ1500(Nikon) equipped with a C11440 camera (Hamamatsu) at a combined magnification of 4 × (1.01μm/pixel). The longitudinal lengths of the worms were measured. Three biological replicates were performed.

#### Survival

Worms were age-synchronized by bleach-based egg isolation followed by starvation in M9 buffer at the L1 stage for at least 24 hours. For all experiments, every genotype was performed in parallel. Synchronized L1 worms were grown to the first day of adulthood, and Day 0 of the lifespan was determined by the onset of egg-laying. During adulthood, worms were transferred to new plates every two days. ~100 animals were assayed for each genotype with 30-40 animals per 6 cm plate. Death was indicated by total cessation of movement in response to gentle mechanical stimulation. Statistical analyses were performed with SPSS (IBM Software) using Kaplan-Meier survival analysis and the log-rank test.

### Stress response assays

#### L1 starvation

Embryos were collected by bleaching gravid adults, resuspended in M9 buffer at a density of 1.5 embryo per microliter, and incubated at 20°C to allow hatching in the absence of food. At intervals of seven days, two aliquots of 30ul were taken from each genotype and counted for survival.

#### Oxidative stress

30 adult worms (D1, 72 hours post arrested L1) per genotype were washed into 24 well plate containing M9 buffer with 2mM hydrogen peroxide, and incubated with gentle shaking at room temperature for 5.5h. Death, defined as absence of movement in liquid buffer upon tapping the plate, was scored immediately.

#### Survival to pathogen

The *E. faecalis* strain OG1RF was grown in BHI broth for 5 hours at 37°C and then spot on 35mm BHI plates. The plates were incubated at 37°C overnight before use.

#### Pseudomonas aeruginosa

PA14 was prepared according to the slow killing protocol. It was cultured in LB medium, seeded onto slow-killing plates, and incubated first at 37°C for 24 hours and then 25°C for 24 hours. *C. elegans* were maintained on *E. coli* OP50 and around 90 animals at L4 stage were transferred onto three technical replicate plates with pathogen for each condition and incubated at 25°C. They were examined for viability every 12 hours for PA14 and every 24 hours for OG1RF. Death was defined as failure to respond to prodding.

#### Cold stress

Animals were allowed to lay eggs for one hour and the progeny synchronized at L4 stage after three days were transferred to 35mm experimental plates seeded with OP50. After 12 hours of growth at 20°C, the experimental plates were placed into a cold chamber that is chilled with ice in a cold room to maintain a plate temperature of 1.5°C. After an interval of time indicated in figures, the animals were allowed to recover at room temperature for 12 hours before prodding with a platinum wire to determine survival. At least 30 animals were used for each condition in a replicate and at least three independent replicates were performed.

#### ER stress

Agar plates were seeded with OP50 *E. coli*, or HT115 for RNAi experiment. ER stress inducer (tunicamycin, Santa Cruz, or dithiothreitol, Sigma) was added to the plate surface resulting in final concentrations as indicated in figures and dried. Synchronized L1 larvae were added to the plates and counted. After three days, numbers of worms at late L4 stage or older (%L4/ adults) were recorded. At least 30 animals were used for each condition in a replicate and at least three independent replicates were performed.

### RNAi knockdown in *C. elegans*

For the RNAi screen, the HT115 bacterial clones targeting different candidates were obtained from both the Vidal and the Ahringer collections. The positive candidates from the screen were sequenced to verify their identity. Specifically, the HT115 RNAi bacterial clones targeting *rsh-1*, *ragc-1, xbp-1, fkb-3, cup-5, nadk-1*, and *nadk-2* were from the Vidal collection, and the ones targeting *vha-8* and *vha-17* were from the Ahringer collection. All the RNAi bacterial clones were grown in LB medium with ampicillin (100μg/mL) for 13 to 14 hours, seeded onto NGM plates with carbenicillin (50μg/mL) and IPTG (1mM), and then incubated at room temperature overnight for inducing the expression of dsRNA. Synchronized L1 were plated onto the bacteria for RNAi knockdown and RNAi screens, whereas RNAi feeding started from the parental generation for *rsh-1* RNAi for consistent results.

Tissue-specific RNAi was achieved using strains with Argonaute mutation *rde-1*(*ne219*) and its tissue specific rescue ^45^. For *cup-5* RNAi experiments, the *cup-*5 RNAi bacteria was diluted two times by L4440 bacteria before seeded onto NGM plates with carbenicilin (50μg/mL) and IPTG (1mM), and then incubated at room temperature overnight for inducing the expression of dsRNA. Synchronized L1 of N2 and *rsh-1* mutants were plated onto the bacteria for *cup-5* RNAi knockdown. Treatment of *cup-5* RNAi bacteria without dilution causes developmental arrest at the first larval stage (L1).

### Tissue-specific activities of UPR

Strains expressing GFP driven by the *hsp-4* (BiP homolog) promoter were grown on OP50 unstressed. Then, L4 animals were transferred to seeded plates containing a sublethal dose of tunicamycin (2μg/mL) for 24h before imaged with fluorescent stereoscope SMZ1500(Nikon) equipped with a C11440 camera (Hamamatsu) at a combined magnification of 18 ×. The head region corresponding to the hypodermis and of the first pair of intestinal cells were measured with ImageJ for median fluorescence intensity and the background fluorescence of the respective image was subtracted for the final value.

### LysoTracker staining in hypodermis and its quantification

LysoTracker DND-26 (Thermo Fisher Scientific) was added at a final concentration of 2μM to the surface of NGM plates pre-seeded with *E. coli*. L4 larvae were transferred to the plate and incubated for 24h before being imaged with confocal microscopy. The hypodermal layer of the body region (the posterior end) was imaged with an 60× objective (PlanAPO N, 1.42N.A., Olympus), scanning along z-axis at 0.6μm per step and the z-axis slices are projected with maximal intensity for analysis. The images were then processed using the following pipeline in CellProfiler ^46^ to identify fluorescent punctae: enhance features (speckles), apply threshold (RobustBackground, global), and identify primary objects (RobustBackground, global). Then the total fluorescent intensity for each puncta was quantified. Different total fluorescent intensity cutoffs for LysoTracker-positive vesicles generate similar results.

### Confocal microscopy

*C. elegans* were anaesthetized in 1% sodium azide in M9 buffer and placed on 2% agarose pad sandwiched between glass microscopic slide and coverslip. The microscope was IX81 (Olympus) connected to an Axiocam ICc3 camera (Zeiss).

### Ultrastructure analysis of hypodermis using transmission electron microscopy

*C. elegans* ultrastructure was imaged following standard Electron Microscopy procedures using a Ted Pella Bio Wave processing microwave with vacuum attachments. The worms were covered in Modified Karnovski’s fixative (2% paraformaldehyde, 2.5% Glutaraldehyde, in 0.1 M Sodium Cacodylate buffer at pH 7.2). While under fixative, *C. elegans* worms tip ends were cut to allow fixative to penetrate the internal tissue. After dissection the dissected worms were incubated for 3 days in the fixative on a rotator. The pre-fixed worms were then microwave and vacuum fixed again, followed by 3× Millipore water rinses, post-fixed with 1% aqueous osmium tetroxide, and rinsed again 3× with Millipore water. Concentrations from 30-100% of Ethanol were used for the initial dehydration series, followed with 3 changes of Propylene oxide as the final dehydrant. Samples were gradually infiltrated with 3 ratios of Propylene oxide and Embed 812, finally going into 3 changes of pure resin under vacuum. Samples were allowed to infiltrate in pure resin overnight on a rotator. The samples were embedded into flat silicone molds and cured in the oven at 62°C for three to five days depending on humidity. The polymerized samples were thin sectioned with a UC7 Ultra-microtome at 48-50 nm and stained with 1% uranyl acetate for thirteen minutes followed by lead citrate for 2.5 minutes before TEM examination. Grids were viewed in a JEOL 1400+ transmission electron microscope at 80kV. Images were captured using an AMT XR-16 mid-mount 16 mega-pixel digital camera. Multi-lamellar bodies were counted for each hypodermal section.

### RSKS-1 phosphorylation analysis using western blot

At least 1,000 worms per genotype arrested at L1 in M9 buffer were plated on seeded NGM plates. They were grown for 45 hours to reach L4 before collected and snap-frozen on dry ice. The samples were lysed in worm lysis buffer (50 mM Tris-HCl pH 7.4, 150 mM NaCl, 1 mM EDTA, 0.1% NP-40) containing a protease and phosphatase inhibitor cocktail (cOmplete™ Protease Inhibitor Cocktail, Cat# 11697498001; PhosSTOP™, Cat# 4906845001; both from Sigma) and homogenized with motorized pellet pestle. The lysates were then centrifuged, and the supernatants were used for protein quantification and western blotting analysis. Next, the proteins were separated with the NuPAGE system (ThermoFisher, 4-12% Bis-Tris protein gel), and transferred to PVDF membrane (ThermoFisher). The membranes were blocked with 5% BSA in TBST. The primary antibody against phosphorylated RSKS-1 is anti-phospho-p70 S6 kinase (Thr389) (1A5) Mouse mAb (Cell Signaling, #9206, 1:1,000), which detects the conserved residue Thr404 in worms and was tested for its specificity using wildtype and *rsks-1* deletion mutant lysates. The anti-beta-actin antibody is from Santa Cruz (sc-47778, 1:2,000). Protein detection was performed using chemiluminescent substrate (ECL™ Western Blotting Reagents, Sigma-Aldrich, GERPN2106) with film exposure (HyBlot CL® Autoradiography Film, Denville Scientific Inc.). Signal intensity was quantified with the Analyze Gels function in ImageJ.

### RNA-seq and data visualization

L1 synchronized worms were plated on OP50 seeded plates after 40 hours of arrest and grown at 20°C. The worms were harvested at mid/ late L4 (after approximately 51h, 54h, 58h, 60h, for N2, *rsh-1(tm6440)*, *rsks-1(ok1255)*, *rsh-1(tm6440);rsks-1(ok1255)*, respectively). Total RNA was extracted from around 2000 worms using Trizol extraction combined with column purification (Qiagen).

Sequencing libraries were prepared using the TruSeq Stranded mRNA Sample Preparation kit (Illumina) following the manufacturer’s instructions. Libraries were pooled together and sequenced using Illumina NextSeq 500 system. Sequencing reads were aligned to C. elegans WS235 genome using Tophat245. Reads were counted using HTseq-count46 and counting data was imported to EdgeR47 for statistical analysis. Statistical significance was defined by adjusted P value (false discovery rate, FDR) of <0.05. A volcano plot was prepared using the MultiplotStudio module on the GenePattern server (https://gp.indiana.edu/gp/).

### Quantification of transcripts with qRT-PCR

RNA was extracted from L4 *C. elegans* with Trizol (Life Technologies) and ethanol precipitation. Total RNA was used to synthesize cDNA with amfiRivert cDNA Synthesis Platinum Master Mix (GenDEPOT). qPCR was performed with KAPA SYBR FAST qPCR Master Mix (2X) Kit (KAPA) in an Eppendorf Realplex 4 PCR machine. Primers used are as follows:

total *xbp-1* FW: ccgatccacctccatcaac, RV: accgtctgctccttcctcaatg
spliced *xbp-1* FW: tgcctttgaatcagcagtgg, RV: accgtctgctccttcctcaatg
*fkb-3* FW: gcaaggattcgacgaagatgga, RV: atccacttggctccgggttt
*fkb-4* FW: ggatttgcgaaagactcgactgg, RV: aggattcgcacggaacagg
*fkb-5* FW: aagaagccatacaccttcaccc, RV: ggtttcctgggatgacgacc
*fkb-7* FW: gccaggacttgataagggtctgaa, RV: tccacaccttgctcttgctc
*fkb-8* FW: cggaatccctaagatgaaagtcgg, RV: agttagggaagcgtttcgagg
*rpl-32* (internal control) FW: agggaattgataaccgtgtccgca, RV: tgtaggactgcatgaggagcatgt

### Mammalian cell lines and culture conditions

Human adipose-derived stem cells (ADSC), obtained from American Type Culture Collection (Manassas, VA, USA) were maintained in MesenPRO RS™ Medium (Thermo Fisher Scientific, Waltham, MA, USA). Human primary mesothelial cells were isolated from omental adipose tissue from consented non-oncological adult donors undergoing elective gastric bypass surgery (Zen-Bio, Inc., Research Triangle Park, NC, USA), and were maintained in mesothelial cell growth medium (MSO-1, Zen-Bio, Inc.). Human cervical adenocarcinoma cell line HeLa from American Type Culture Collection was maintained in DMEM medium supplemented with 10% fetal bovine serum, 2 mM glutamine and penicillin-streptomycin (Life Technologies Corp., Carlsbad, CA, USA). All cells were cultured in humidified incubator with 5% CO2 at 37 °C. HeLa cells were tested negative for mycoplasma contamination and were authenticated by short tandem repeat profiling in the Characterized Cell Line Core Facility at MD Anderson Cancer Center.

### HDDC3 localization with immunofluorescence microscopy

Cells cultured on glass coverslips were fixed with 4% paraformaldehyde and blocked with 1% bovine serum albumin (BSA) and 0.1% saponin. The fixed cells were then incubated with antibodies diluted in 1% BSA and 0.1% saponin at ambient temperature: 2 h with primary antibodies (anti-HDDC3, 1:25, cat# HPA040895, Sigma-Aldrich Co., St. Louis, MO, USA; and anti-LAMP2, 1:200, cat# sc-18822, Santa Cruz) followed by 1 h with secondary antibodies (Alexa Fluor 568 anti-mouse, 1:100, A-11031, Thermo Fisher Scientific; and Alexa Fluor 488 anti-rabbit, 1:100, A-11034, Thermo Fisher Scientific). Next, the cells were mounted in antifade mounting medium with 4′,6-diamidino-2-phenylindole (DAPI; Thermo Fisher Scientific) and visualized with IX81 confocal microscope (Olympus) connected to an Axiocam ICc3 camera (Zeiss).

### Mammalian S6K phosphorylation by western blot

HDDC3 was transiently silenced in HeLa cells by transfecting MISSION predesigned siRNAs (SASI_Hs01_00101056 and SASI_Hs01_00101057, Sigma-Aldrich Co.) duplexed with Lipofectamine RNAiMAX (Life Technologies Corp.) at a final concentration of 20 nM for 8 h. MISSION siRNA universal negative control #1 (Sigma-Aldrich Co.) was used as negative control. Then, cell extracts were prepared in a RIPA buffer (20 mM sodium phosphate, 150 mM NaCl pH 7.4, 1% Nonidet P-40, 0.1% SDS, and 0.5% deoxycholic acid) containing a protease and phosphatase inhibitor cocktail (cOmplete™ Protease Inhibitor Cocktail and PhosSTOP™ from Sigma). Proteins were separated on SDS-polyacrylamide gels and electrophoretically transferred to an Immobilon polyvinylidene fluoride membrane (EMD Millipore, Billerica, MA, USA). The membranes were incubated with primary antibodies overnight at 4⁰C, and then were incubated with appropriate horseradish peroxidase-conjugated secondary antibodies at 1:10,000 dilution (Thermo Fisher Scientific) for 1 h at ambient temperature. Signals were developed using ECL chemiluminescence detection reagents (Denville Scientific Inc., Holliston, MA, USA) and visualized on X-ray film (Fujifilm, Tokyo, Japan). The following primary antibodies were used: anti-p-S6K (T389) (1:1000, cat#9234, Cell Signaling Technology, Beverly, MA, USA), anti-S6K (1:1000, cat# 9202, Cell Signaling Technology), and β-actin (1:5,000; clone AC-15, Sigma-Aldrich Co.).

### Modeling of *C. elegans* RSH-NADPH complex

The *C. elegans* RSH protein (ZK909.3) sequence was extracted from wormbase (https://wormbase.org/). To build accurate homology models, two ligand free crystal structures of SpoT family proteins, *H. sapiens* RSH (PDB ID: 3NR1) and *D. melanogaster* RSH (PDB ID: 3NQW) were used as multiple templates to generate initial *C. elegans* RSH (CeRSH) homology model in the software Modeller ^47^. Then the crystal structure of *H. sapiens* RSH that was solved in the presence of a NADPH co-factor (PDB ID: 5VXA) was used as a template for Modeller to extract NADPH and satisfy distance restrains into CeRSH models. The model with the lowest DOPE global score was selected as the final *C. elegans* RSH model. The structure images were generated by UCSF Chimera ^48^.

### Statistics and reproducibility

Sample size was determined based on experience, variability of the results and previous literature using the same assays. All experiments have been repeated with a minimum of three biologically independent replicates. For randomization, *C. elegans* were mixed well before allocating to different conditions. Individuals of the same age across conditions were picked randomly for imaging.

Statistical analyses used GraphPad software. In experiments comparing difference with genotypes and dose, two-way ANOVA was performed followed by post hoc analysis. Two-sided unpaired Student’s *t*-test was used for pairwise comparison and one-way ANOVA or multiple *t*-test for comparing multiple groups. Normality of data is assumed for the other experiments. Data of bar charts are represented as mean ± standard error of the mean (S.E.M), and data of dot plots as mean ± standard deviation (SD). The provided *n* values in the dot plots refer to sample sizes and those in the legend to number of independent replicates. \*\*\**p<0.001*, \*\**p<0.01*, \**p<0.05*, n.s. *p>0.05*.

## Data availability

The data that support the findings of this study are available from corresponding authors upon request.

## Acknowledgements

We thank J. Mello (Harvard Medical School, USA) for providing *rde-1* rescue strains, J. Mamrosh (Caltech, USA) for SJ4005, Xiaochen Wang for XW5735 strain (Chinese Academy of Science, China), M. Cruz from D. Garsin lab (UTHealth) for preparing PA14 and *E. faecalis*, National BioResource Project for *rsh-1(tm6440)*, Caenorhabditis Genetics Center for mutant and transgenic strains, and wormbase for valuable *C. elegans* information. The work was supported by NIH grants R01AG045183 (M.C.W.), R01AT009050 (M.C.W.), DP1DK113644 (M.C.W.), NIH grant R35GM127088 (J.D.W.), and by HHMI faculty scholar (J.D.W.) and HHMI investigator support (M.C.W.).

## Author Contributions

K.H.M and M.C.W wrote the manuscript. K.H.M, M.C.W and J.D.W conceived and designed the study. K.H.M and Q.Z. conducted the experiments and analyzed the results. Q.Z. conducted structural simulation. L.D, K.H.M and P.H prepared and analyzed the TEM samples. K.H.M and P.H prepared and analyzed the RNA-seq data. C.A. performed mammalian cell experiments. J.Y performed and analyzed the ppGpp hydrolysis experiment. Y.Y generated the *col-12p::lmp-1::rfp::3xha* strain and optimized lysosome immunopurification protocol. M.A. and D.S. consulted with the optimization of lysosomes immunopurification protocol.

