## Supplementary figures and tables for "Lysosomal nucleotide metabolism regulates ER proteostasis through mTOR signaling"

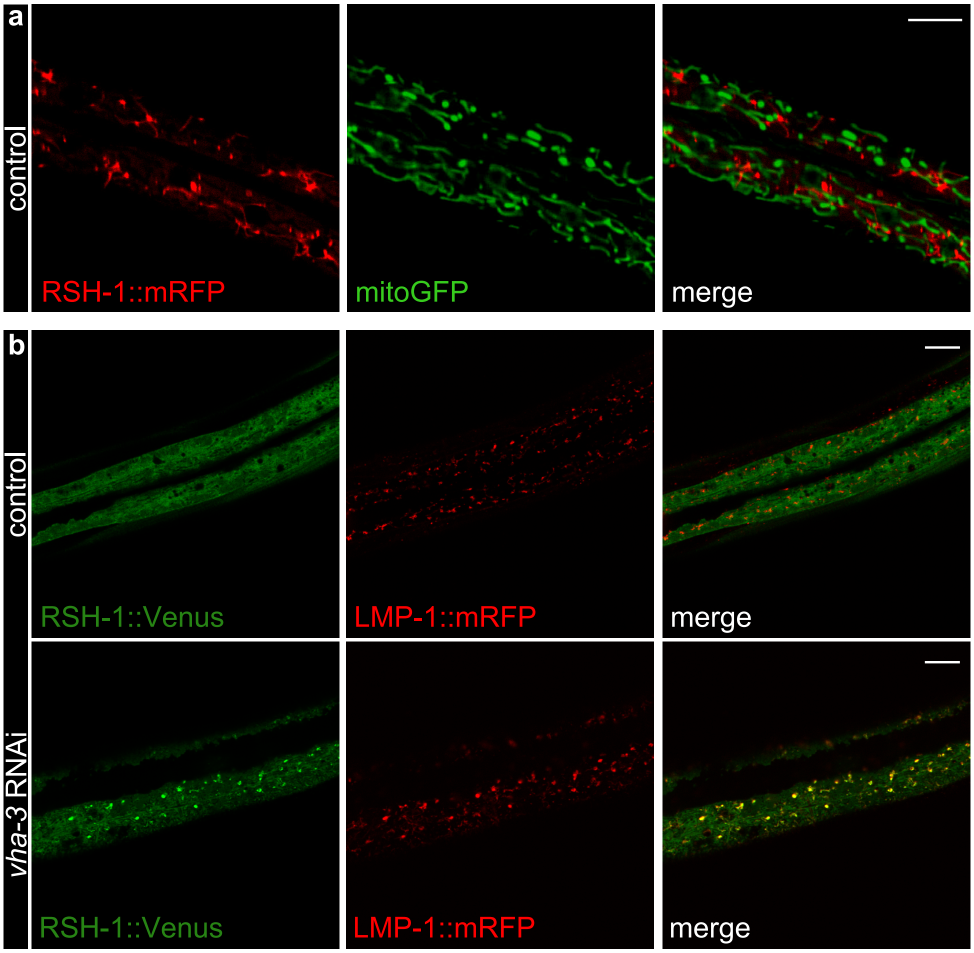

**Extended Data Figure 1. Subcellular localization of fluorescent protein fusions of RSH-1 in hypodermal cells**

**a**) RSH-1::mRFP does not overlap with mitoGFP signals in hypodermal cells.

**b**) RSH-1::Venus overlaps with LMP-1::mRFP in hypodermal cells upon *vha-3* RNAi knockdown.

Scale bar = 10μm.

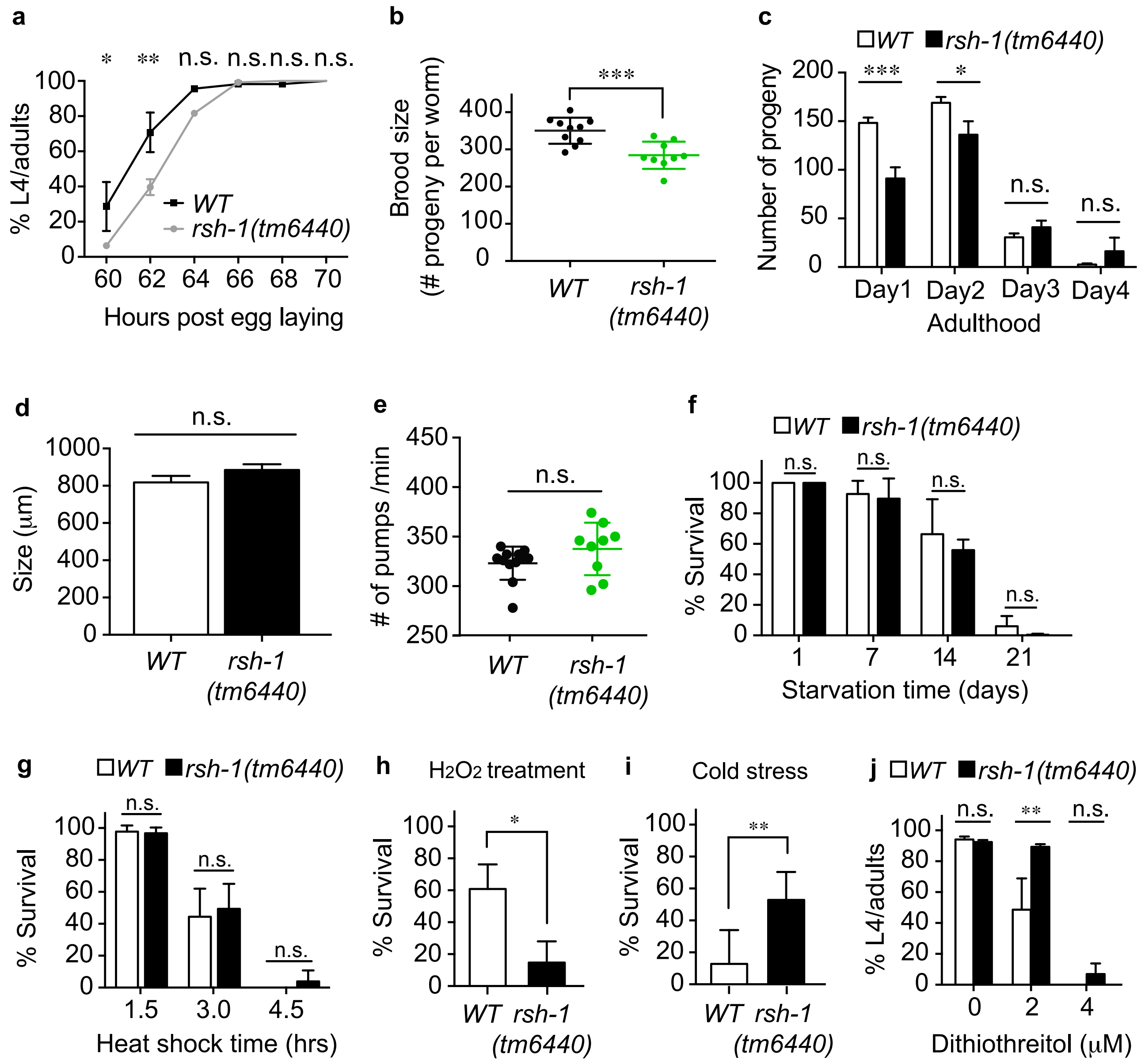

**Extended Data Figure 2. Physiological activities and stress responses in the *rsh-1* mutants**

**a - e**) Under an unstressed condition, the *rsh-1(tm6440)* mutants have slightly delayed development (**a**) and reduced brood size (**b** and **c**) (*n* = 10 for WT, *n* = 9 for *rsh-1(tm6440)*), but are similar to WT in their body length (**d**) (*n* = 3 replicates) and food intake rate (**e**) (*n* = 24 for WT, *n* = 19 for *rsh-1(tm6440)*).

**f** and **g**) The *rsh-1(tm6440)* mutant shows no significant difference in survival under starvation (**f**, *n*=3 replicates), or heat shock (**g**, *n*=3 replicates), when compared to WT.

**h**) The *rsh-1(tm6440)* mutant is more sensitive to oxidative stress induced by hydrogen peroxide than WT (*n* = 3 replicates).

**i** and **j**) The *rsh-1(tm6440)* mutant shows increased resistance to cold shock (**i**, $n = 4$ replicates) and to dithiothreitol-induced ER stress (**j**, $n = 3$ replicates), when compared to WT.

Error bars represent SEM. ***$p<0.001$, **$p<0.01$, *$p<0.05$, n.s. $p>0.05$ by two-way ANOVA (a, c, f, g, j) or *t*-test (b, d, e, h, i).

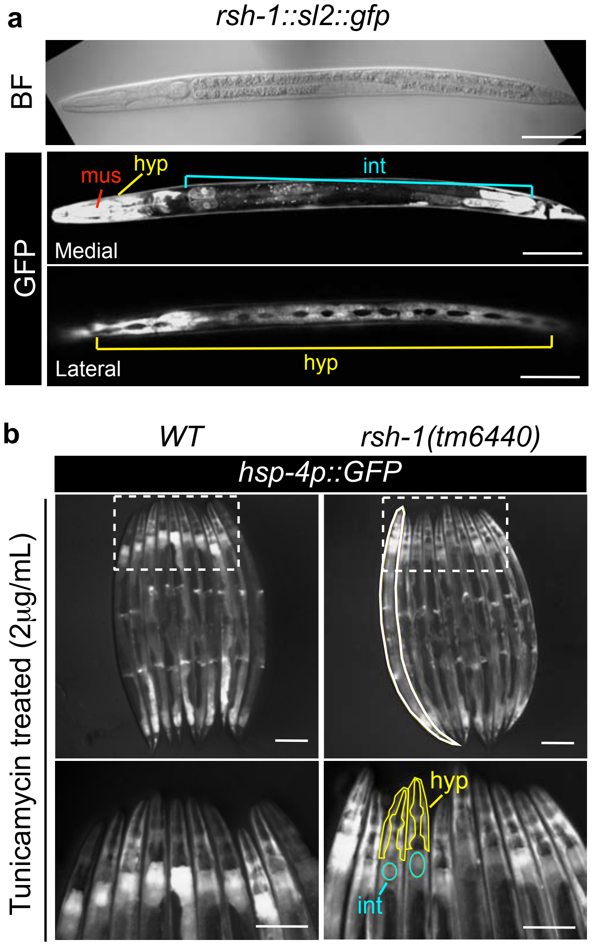

**Extended Data Figure 3. RSH-1 functions in the hypodermis to regulate UPR$^{ER}$.**

**a**) A GFP reporter driven by the *rsh-1* genomic regulatory element shows *rsh-1* expression in pharyngeal muscle (mus), hypodermis (hyp) and intestine (int). The representative images (BF, bright field) are from a L3 larva. Scale bar = 50μm.

**b**) Upon tunicamycin treatments, the level of the *hsp-4* (ER chaperone) GFP reporter is increased in both hypodermal and intestinal cells, and the hsp-4 induction in hypodermal cells is specifically enhanced in the *rsh-1(tm6440)* mutant compared to WT. The high-magnification images of the dashed boxed areas are shown in lower panels, and solid yellow and cyan outlines indicate hypodermal and intestinal regions used for signal quantification in Fig. 3e. Scale bar = 200μm.

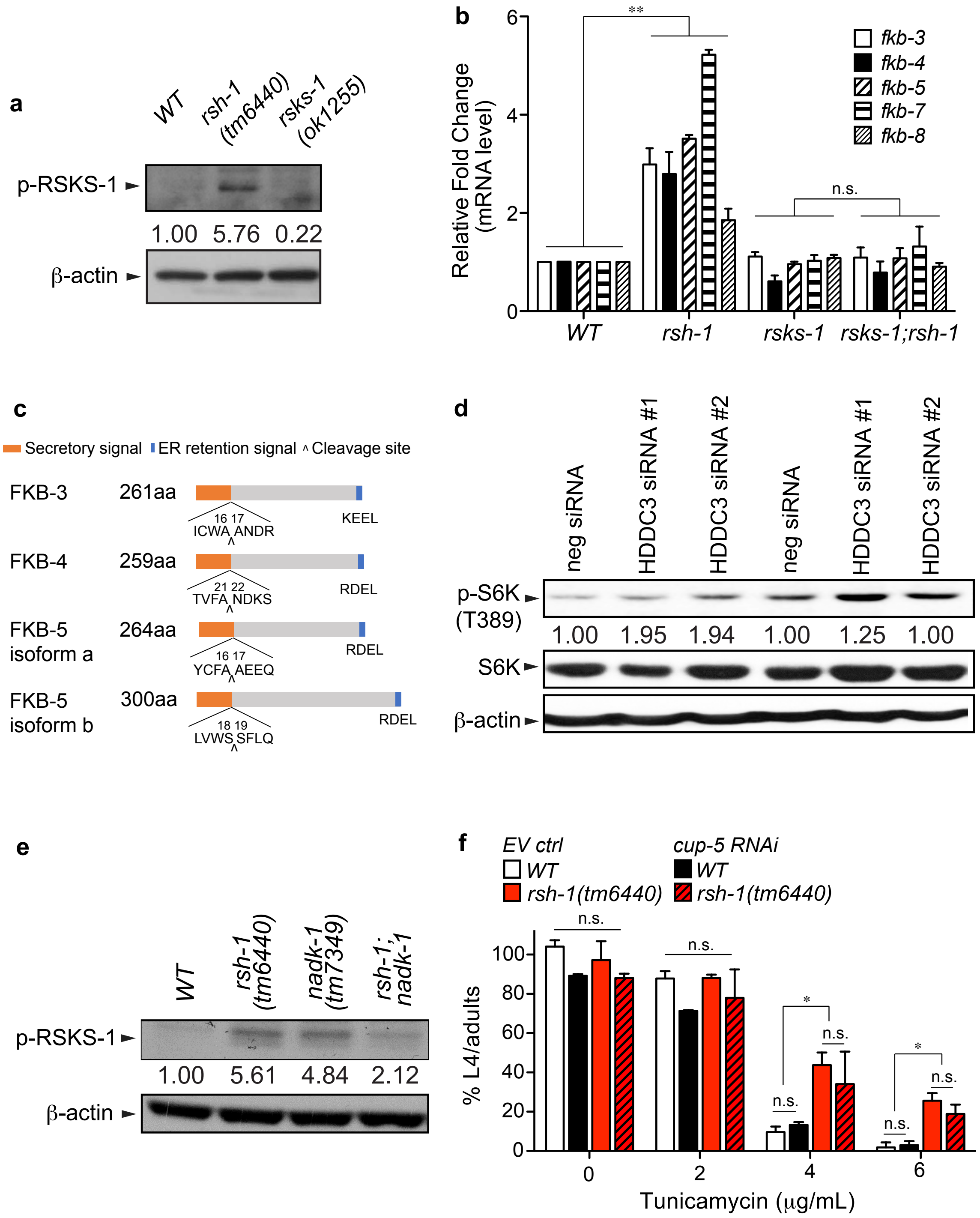

**Extended Data Figure 4. Regulation of mTOR and ER by RSH-1**

**a**) Immunoblotting shows an increased level of phosphorylated RSKS-1(Thr404) in the unstressed *rsh-1(tm6440)* mutants. The values below the blot indicate the relative level of phosphorylated RSKS-1 after normalization to β-actin. Replicate #2.

**b**) qRT-PCR shows that the transcriptional levels of *fkb-3,4,5,7,* and *8* are up-regulated in the *rsh-1(tm6440)* mutants, and these inductions are fully suppressed by the *rsks-1(ok1255)* mutant. Error bars represent SEM. *n* = 3 replicates. **$p<0.01$, n.s. $p>0.05$ by two-way ANOVA.

**c**) FKB-3, FKB-4 and FBK-5 have a secretory signal peptide and an ER retention signal at the N- and C- termini, respectively. Locations of the secretory signal peptide and the cleavage site are predicted by SignalP3.0 [49].

**d**) The level of phosphorylated p70 S6K is increased in HeLa cells after transfection with HDDC3-specific siRNAs, compared to control siRNA. Relative p-S6K levels normalized to the corresponding total p70 S6 kinase level are indicated. Replicate #2 and #3.

**e**) The *rsh-1(tm6440)* and the *nadk-1 (tm7349)* mutants both have increased RSKS-1 phosphorylation, compared to WT. The induction is abolished in the double mutant. Replicate #2.

**f**) RNAi inactivation of *cup-5*, the *C.elegans* ortholog of human lysosomal calcium channel MCOLN1, does not change the ER stress tolerance of the *rsh-1(tm6440)* mutants. Error bars represent SEM. n = 3 replicates for each group.

**Supplementary Table 1 Summary of lysosomal genes screened by RNAi**

**Supplementary Table 2 Summary of nucleotide metabolic genes screened by RNAi**

**Supplementary Table 1. Summary of lysosomal genes screened by RNAi**

| Category | Sequence name | Gene name | Suppression of ER tolerance in rsh-1 | Reduction of ER tolerance in WT |
|---|---|---|---|---|
| lysosome | F49C12.13 | *vha-17* | Yes | Yes |
| lysosome | D2096.3 | *aagr-1* | Yes | No |
| lysosome | C17H12.14 | *vha-8* | No | Yes |
| lysosome | Y38F2AL.4 | *vha-3* | No | Yes |
| lysosome | T14F9.1 | *vha-15* | Severe delay on *WT* and *rsh-1* w/o TM | |
| lysosome | F46F11.5 | *vha-10* | Severe delay on *WT* and *rsh-1* w/o TM | |
| lysosome | Y38F2AL.3 | *vha-11* | Severe delay on *WT* and *rsh-1* w/o TM | |
| lysosome | F20B6.2 | *vha-12* | Severe delay on *WT* and *rsh-1* w/o TM | |
| lysosome | T01H3.1 | *vha-4* | Severe delay on *WT* and *rsh-1* w/o TM | |
| lysosome | R10E11.8 | *vha-1* | Severe delay on *WT* and *rsh-1* w/o TM | |
| lysosome | F35H10.4 | *vha-5* | No | |
| lysosome | ZK484.2 | *haf-9* | No | |
| lysosome | K11D2.2 | *asah-1* | No | |
| lysosome | C33C12.8 | *gba-2* | No | |
| lysosome | W06B4.3 | *vps-18* | No | |
| lysosome | W07B8.5 | *cpr-5* | No | |
| lysosome | ZC190.1 | *cln-3.3* | No | |
| lysosome | F35E8.11 | *cdr-1* | No | |
| lysosome | F38A6.3 | *hif-1* | No | |
| lysosome | C03B1.12 | *lmp-1* | No | |
| lysosome | C07B5.5 | *nuc-1* | No | |
| lysosome | F52E1.10 | *vha-18* | No | |
| lysosome | F55H2.2 | *vha-14* | No | |
| lysosome | Y110A7A.12 | *spe-5* | No | |
| lysosome | Y49A3A.2 | *vha-13* | No | |
| lysosome | C30F8.2 | *vha-16* | No | |
| lysosome | Y55H10A.1 | *vha-19* | No | |
| lysosome | VW02B12L.1 | *vha-6* | No | |
| lysosome | ZK637.8 | *unc-32* | No | |
| lysosome | F53F10.4 | *unc-108* | No | |

| | | | |
|---|---|---|---|
| lysosome | C01G8.2 | *cln-3.2* | No |
| lysosome | T27A1.5 | - | No |
| lysosome | C33C12.3 | *gba-1* | No |
| lysosome | R05F9.12 | *aagr-2* | No |
| lysosome | F59G1.3 | *vps-35* | No |
| lysosome | C32D5.9 | *lgg-1* | No |
| lysosome | C56C10.1 | *vps-33.2* | No |
| lysosome | B0252.2 | *asm-1* | No |
| lysosome | C41C4.7 | *ctns-1* | No |
| lysosome | R06F6.2 | *vps-11* | No |
| lysosome | F37H8.5 | - | No |
| lysosome | W03C9.3 | *rab-7* | No |
| lysosome | K09E4.4 | - | No |
| lysosome | R13A5.1 | *cup-5* | No |
| lysosome | B0303.9 | *vps-33.1* | No |
| lysosome | Y37D8A.2 | - | No |
| lysosome | Y43F4B.7 | - | No |
| lysosome | T19E7.3 | *bec-1* | No |
| lysosome | C28C12.7 | *spp-10* | No |
| lysosome | T20D3.7 | *vps-26* | No |
| lysosome | F07B10.1 | *cln-3.1* | No |
| lysosome | F11E6.1 | *gba-3* | No |
| lysosome | F41E6.6 | *tag-196* | No |
| lysosome | F41E6.13 | *atg-18* | No |
| lysosome | T19B10.3 | - | No |
| lysosome | C42C1.4 | - | No |
| lysosome | F57F5.1 | - | No |
| lysosome | C52E4.1 | *cpr-1* | No |
| lysosome | R07B7.11 | *gana-1* | No |
| lysosome | T03E6.7 | *cpl-1* | No |
| lysosome | F52D1.1 | *aagr-4* | No |
| lysosome | C05D9.2 | *lmp-2* | No |
| lysosome | T14F9.3 | *hex-1* | No |

| | | | |
|---|---|---|---|
| lysosome | K09C4.8 | *sul-1* | No |
| lysosome | F14H12.4 | *cst-1* | No |
| lysosome | F02E8.6 | *ncr-1* | No |
| lysosome | C25B8.3 | *cpr-6* | No |
| lysosome | H22K11.1 | *asp-3* | No |
| lysosome | C54D2.4 | *sul-3* | No |
| lysosome | R09F10.1 | - | No |
| lysosome | ZK721.1 | *chup-1* | No |
| lysosome | R12H7.2 | *asp-4* | No |
| lysosome | F09B12.3 | - | No |
| lysosome | Y111B2A.8 | *aakg-1* | No |
| lysosome | Y24F12A.2 | *ragc-1* | No |
| lysosome | Y105E8B.9 | - | No |
| lysosome | Y57G11C.13 | *arl-8* | No |
| lysosome | Y39B6A.20 | *asp-1* | No |
| lysosome | F41C3.5 | - | No |
| lysosome | T20G5.10 | *blos-1* | No |
| lysosome | Y39A1A.1 | *epg-6* | No |
| lysosome | T23D8.2 | *tsp-7* | No |
| lysosome | Y4C6B.6 | *gba-4* | No |
| lysosome | F27E5.1 | - | No |
| lysosome | D1014.1 | *sul-2* | No |
| lysosome | R02E12.6 | *hrg-1* | No |
| endosome | K02D10.5 | *snap-29* | Severe delay on *WT* and *rsh-1* w/o TM |
| endosome | R07G3.1 | *cdc-42* | No |
| endosome | F25B3.1 | *ehbp-1* | No |
| endosome | C06A6.3 | *mvb-12* | No |
| endosome | R02E12.6 | *hrg-1* | No |
| endosome | F35H10.4 | *vha-5* | No |
| endosome | D1037.4 | *rab-8* | No |
| endosome | F53F10.4 | *unc-108* | No |
| endosome | C34G6.7 | *stam-1* | No |
| endosome | T23H2.5 | *rab-10* | No |

| | | | |
|---|---|---|---|
| endosome | F18C12.2 | *rme-8* | No |
| endosome | W06D4.5 | *snx-3* | No |
| endosome | F26H9.6 | *rab-5* | No |
| endosome | F25D7.1 | *cup-2* | No |
| endosome | ZK1248.10 | *tbc-2* | No |
| endosome | C56C10.3 | *vps-32.1* | No |
| endosome | W03C9.3 | *rab-7* | No |
| endosome | R10E11.3 | *usp-46* | No |
| endosome | C38H2.1 | *tbc-8* | No |
| endosome | Y39A1A.5 | *rabx-5* | No |
| endosome | C18D11.2 | *maa-1* | No |
| endosome | C09G12.9 | *tsg-101* | No |
| endosome | T11F8.3 | *rme-2* | No |
| endosome | F58G6.1 | *amph-1* | No |
| endosome | CD4.4 | *vps-37* | No |
| endosome | Y32F6B.3 | *crp-1* | No |
| endosome | T10G3.5 | *eea-1* | No |
| endosome | F49E7.1 | *rme-6* | No |
| endosome | C05D9.1 | *snx-1* | No |
| endosome | F02E8.6 | *ncr-1* | No |
| endosome | F14B8.2 | *sid-5* | No |
| endosome | F45E1.7 | *sdpn-1* | No |
| endosome | ZK721.1 | *chup-1* | No |
| endosome | K09A9.2 | *rab-14* | No |
| endosome | Y87G2A.10 | *vps-28* | No |
| endosome | R10E12.1 | *alx-1* | No |
| endosome | Y49E10.11 | *tat-1* | No |
| autophagy | T19E7.3 | *bec-1* | No |
| autophagy | F38A6.1 | *pha-4* | No |
| autophagy | K07A1.2 | *dut-1* | No |
| autophagy | T23G11.7 | - | No |
| autophagy | Y106G6A.2 | *epg-8* | No |
| autophagy | M01E5.6 | *sepa-1* | No |

| | | | |
|---|---|---|---|
| autophagy | C32D5.9 | *lgg-1* | No |
| autophagy | D2085.2 | *atg-10* | No |
| autophagy | W03C9.3 | *rab-7* | No |
| autophagy | B0336.8 | *lgg-3* | No |
| autophagy | T26A5.9 | *dlc-1* | No |
| autophagy | ZK593.6 | *lgg-2* | No |
| autophagy | M7.5 | *atg-7* | No |
| autophagy | C10F3.5 | *pcm-1* | No |
| autophagy | K04A8.5 | *lipl-4* | No |
| autophagy | F41E6.13 | *atg-18* | No |
| autophagy | Y60A3A.1 | *unc-51* | No |
| autophagy | Y39A1A.1 | *epg-6* | No |
| autophagy | M03A8.2 | *atg-2* | No |

**Supplementary Table 2. Summary of nucleotide metabolic genes screened by RNAi**

| Sequence name | Gene name | Suppression of ER tolerance in rsh-1 | Reduction of ER tolerance in WT |
|---|---|---|---|
| Y77E11A.2 | *nadk-1* | Yes | No |
| R151.2 | - | Yes | No |
| Y17G7B.10 | *nadk-2* | Yes | No |
| F46H6.2 | *dgk-2* | No | |
| F42A9.1 | *dgk-4* | No | |
| K06A1.6 | *dgk-5* | No | |
| Y105E8B.5 | *hprt-1* | No | |
| T19B4.3 | - | No | |
| T10B11.2 | *cerk-1* | No | |
| R07H5.8 | *adk-1* | No | |
| B0228.7 | - | No | |
| C37H5.6 | *adss-1* | No | |
| M106.4 | *gmps-1* | No | |
| T22D1.3 | - | No | |
